# A novel mechanism for the uptake of enterobactin-chelated ferric ions by the mitochondria and subsequent reduction to ferrous ions inside

**DOI:** 10.1101/2024.09.05.611381

**Authors:** Abeg Dutta, Anutthaman Parthasarathy, Namasivayam Ganesh Pandian

## Abstract

Conventional knowledge of mitochondrial iron metabolism talks about the export of iron in its doubly charged state once it is reduced inside the cell until it reaches the mitochondria. At physiological oxygen tension and pH 7.4, the comparatively soluble Fe(II) is easily available. Fe(III) hydrolyses to generate insoluble ferric hydroxides. Iron must be regularly chaperoned because of its near insolubility and potential toxicity due to redox activity. All tissues pick up iron through the binding of transferrin (Tf) to the transferrin receptor 1 (TfR1), followed by the complex’s internalisation through receptor-mediated endocytosis. The low pH created by the operation of a proton pump within the endosome reduces Tf’s affinity for iron. Importantly, the TfR1 promotes iron escape from Tf in the pH range (pH 5-5.5) reached by the endosome. A "trap," such as pyrophosphate, is needed *in vitro* for iron release from Tf. However, a physiological chelator that can play this role has not yet been discovered. Fe(III) is hypothesised to be reduced to Fe(II) in erythroid cells by a ferrireductase known as the six-transmembrane epithelial antigen of the prostate 3 in the endosomal membrane after being released from Tf in the endosome. Following this, the divalent metal transporter-1 (DMT1) transports Fe(II) through the endosomal membrane and, it is generally accepted that this generates the cytosolic labile or chelatable iron pool. This reservoir of iron is believed to provide the metal for metabolic requirements, including as iron intake by the mitochondrion for haem and ISC synthesis, as well as storage in the cytosolic protein ferritin. The possibility of Fe(III) entering the mitochondria has not been explored before. A series of dockings shows that binding of the alpha subunit to a complex of Fe(III) with enterobactin is as stable as the binding with of the same chelator with Fe(II). Enterobactin is a bacterial iron chelator thought to play an integral role in host iron metabolism. Our results suggest an interesting possibility of iron being trafficked to the mitochondria as Fe(III). We propose a potential mechanism of Fe(III) trafficking and subsequent reduction to Fe(II) inside the mitochondria.

## Introduction

### Iron in the mitochondria

The mitochondrion is primarily valued for its function in energy transfer. Less well recognised is the fact that this organelle serves as a focal point for the metabolism of iron, the most prevalent transition metal in cells. The ability of the mitochondrion to catalyse electron transport via haem- and iron sulphur cluster (ISC)-containing proteins and utilise this procedure in energy transduction is made possible by iron’s reversible oxidation states. Given this fact alone, it is understandable why the mitochondrion is so important for iron metabolism. It is the only location where haem is synthesised and is a significant source of ISCs (found in both mitochondria and cytoplasm). Interventions such as the removal of mitochondrial iron by the small molecule ironomycin result in triggering cell death (1). Indeed, frataxin (FXN) is a small acidic protein containing intrinsically disordered regions, which regulates iron traffic to the ISC machinery, and is a key regulator of iron homeostasis (2). FXN is imported into the mitochondria, can bind both Fe(II) and Fe(III), and is a regulator of the iron-dependent apoptosis or ferroptosis discovered in 2012 (3), which has been linked to several diseases (4).

Due to its unmatched versatility as a biological catalyst, iron is a crucial nutrient. Iron is therefore vital for growth, but it is difficult for organisms to acquire due to a paradoxical situation created by the same chemical features which also allow for its adaptability. This is because the relatively soluble Fe(II) is easily converted to Fe(III) under physiological oxygen tension, which upon hydrolysis results in insoluble ferric hydroxides. Iron must be regularly chaperoned because of its near insolubility and probable toxicity due to redox activity. In fact, to meet the organism’s iron needs, specialised molecules for the acquisition, transportation, and storage of iron in a soluble, harmless form have evolved (5).

### Current models of mitochondrial iron transport

Differential transferrin (Tf) is especially responsible for transporting iron in the blood under physiologically normal circumstances. Tf binds to the transferrin receptor (TfR1) to provide iron to all tissues, which are subsequently internally processed by receptor-mediated endocytosis. The endosome’s low pH, brought about by proton pump action, reduces Tf’s affinity for iron. The divalent metal transporter-1 (DMT1) is responsible for moving iron from the endosome and into the chelatable or labile iron pool. It is the mitochondrial inner membrane that contains proteins suitable for importing Fe(II) into the mitochondrial matrix, although the mechanisms are not well understood yet.

As soon as iron enters the mitochondrion, it can be used to make haem, ISC, or be stored as mitochondrial ferritin. As mitochondria are a primary source of cytotoxic Reactive Oxygen Species (ROS), it is crucial that mitochondrial iron is kept in a stable form to restrict oxidative damage. Therefore, it is likely that iron is transported within the mitochondrion subject to strict regulation, like how it is transported in the cytosol, in a form remote from the aqueous environment within the hydrophobic pockets of interacting proteins that constitute iron transport channels (5).

### Proposal for an alternative model

Qi and Han (2018) demonstrated that enterobactin, the bacterial siderophore, promotes growth and the labile iron pool in *C. elegans*. It does so by binding to the alpha subunit of the mitochondrial ATPase and further demonstrated the consistency of this mechanism in mammalian cells. This happens inside the mitochondria and has been shown to be operational independently of ATP synthase. Originally presumed to have a negative effect on host iron uptake, these results suggest a novel commensal-host interaction mechanism mediated by enterobactin for mitochondrial iron uptake, especially given the high prevalence of enterobactin-producing bacteria in the human gut (6).

## Results

### Initial docking results

The first phase of docking involves comparing the binding affinities of Fe(II) and Fe(III) to the enterobactin-alpha subunit complex. The human mitochondrial alpha subunit of the F1 ATPase has been retrieved from AlphaFold2 (P25705-ATPA_HUMAN). AlphaFold2 is the latest tool which is bringing the goal of accurate structure prediction from the primary sequence (at resolutions similar to experimentally solved structures) closer (7). Autodock Vina (8, 9) was used initially for the docking of the enterobactin-iron-alpha via four different approaches. The first two approaches involve docking of Fe(II), with the first approach involved in docking of Fe(II) to the enterobactin-alpha subunit docked complex and the second approach involving the docking of Fe(II) with enterobactin first followed by docking to the alpha subunit. The same then has been done for Fe(III) to constitute approaches 3 and 4. Approaches 2 and 4 have been replicated into a and b as the docking of Fe(II) and Fe(III), respectively to enterobactin yielded a set of docked sites out of which the second most stable confirmation happens to be one available in Pubchem (ID: 16048613), instead of the one yielded by Autodock Vina. Hence, docking was performed for the theoretically obtained most stable conformation obtained, in addition to the second most stable confirmation of iron-enterobactin. For each of the approaches, interactive protein-ligand diagrams have been obtained in LIGPLOT+ (10) and PyMol (11). The different docking approaches are summarised in **Table 1**.

**Table 1:** The various docking approaches used in this study.

| Approach | First docking | Second docking |
| --- | --- | --- |
| 1 | Ent-alpha | Fe2+-(Ent-alpha) |
| 2a | Fe2+-Ent | Alpha-(Fe2+-Ent), most stable confirmation from first docking |
| 2b | Fe2+-Ent | Alpha-(Fe2+-Ent), second most stable confirmation from first docking |
| 3 | Ent-alpha | Fe3+-(Ent-alpha) |
| 4a | Fe3+-Ent | Alpha-(Fe3+-Ent), most stable confirmation from first docking |
| 4b | Fe3+-Ent | Alpha-(Fe3+-Ent), second most stable confirmation from first docking |

Table 2 represents all the binding affinities obtained from the docking approaches outlined in **Table 1**. The most negative binding affinities have been observed for Fe(II) when, docked to enterobactin, which is in turn docked to the alpha subunit of ATP synthase (for the most stable conformation of the Fe(II)- enterobactin docked states), as observed in **Table 2** for Approach 2a. For Fe(III), it is when the Fe(III) ion has been docked to the enterobactin-alpha complex (enterobactin docked to alpha subunit), as observed in **Table 2** for Approach 3. Both have binding affinity of -9.4 kcal/mol. It is possible that the Fe(II)-enterobactin-alpha complex could be more or less as stable as the complex formed by Fe(III). The *in silico* docking parameters used in this study do not yet account for the lipophilic and acidic nature of the mitochondrial matrix (**Fig S1**). The in silico docking parameters do not account for the lipophilic and acidic nature of the mitochondrial matrix. **Fig S2** shows all the binding affinities obtained from molecular docking, with the most negative values at Approaches 2a and 3 and the least negative for Approach 2b for the least stable set of conformations. Although mitochondrial iron uptake and transport has been mostly studied in the context of Fe(II) uptake, it is possible that enterobactin facilitates uptake of Fe(III) in the mitochondria. Once in the mitochondrial matrix, a ferric reductase can reduce the Fe(III) to Fe(II) without the requirement of ATP.

**Table 2:** The values of binding affinities (in kcal/mol) obtained from Autodock Vina for the approaches outlines above in Table 1. The primary focus is on the most negative values.

| S. No | Approach 1 | Approach 2a | Approach 2b | Approach 3 | Approach 4a | Approach 4b |
| --- | --- | --- | --- | --- | --- | --- |
| 1 | -8.9 | -9.4 | -8.7 | -9.4 | -8.6 | -8.4 |
| 2 | -8.8 | -8.9 | -8.3 | -8.9 | -8.5 | -8.4 |
| 3 | -8.6 | -8.7 | -8.2 | -8.3 | -8.2 | -8.4 |
| 4 | -8.4 | -8.5 | -7.8 | -8.2 | -8.1 | -8.3 |
| 5 | -8.3 | -8.4 | -7.7 | -8.2 | -7.8 | -8.2 |
| 6 | -8.2 | -8.2 | -7.6 | -8.2 | -7.7 | -8.1 |
| 7 | -8.2 | -8.1 | -7.4 | -8.1 | -7.7 | -8.1 |
| 8 | -8.1 | -8.1 | -7.3 | -8.0 | -7.6 | -8.0 |
| 9 | -7.9 | -8.1 | -7.2 | -8.0 | -7.6 | -8.0 |

**Figures S3-S12** represent the docking snapshots obtained from Autodock Vina and 2D protein interaction diagrams from LigPLot+. **Fig S3** shows the most and second most stable conformations obtained for Approach 1. **Fig S7** shows the interaction of enterobactin with the Ser184, Asp390 and Arg416 of the chain A of the alpha subunit of the mitochondrial ATPase by hydrogen bonds, detailing the docking snapshot in **Fig S3**. Non ionic interactions are denoted by red arcs with outward lines denoting the direction of the interaction/s.

### Docking (Part 2)

The binding of enterobactin to the alpha subunit produces a very stable complex and this stability is more than the iron-enterobactin complexes *in silico* as can be seen in **Table 3**. We hypothesise that this demonstrates the possibility of a Fe(II)/(III)-Enterobactin complex entering the mitochondria and undergoing dissociation to yield free iron and a more stable Enterobactin-alpha subunit complex. This subject has been explored in detail in the discussion section.

**Table 3:** Separate binding affinities for the Fe-enterobactin and enterobactin-alpha subunit complexes.

| Complex | Binding affinity (kcal/mol) |
| --- | --- |
| Enterobactin-alpha subunit | -8.7 |
| Fe(II)-enterobactin (approach 2a) | -0.9 |
| Fe(II)-enterobactin (approach 2b) | -0.7 |
| Fe(III)-enterobactin (approach 4a) | -0.9 |
| Fe(III)-enterobactin (approach 4b) | -0.7 |

### Docking (Part 3)

A set of four molecular docking algorithms were carried out in Autodock Vina for Fe(II) and Fe(III) each with Bovine mitochondrial ATPase and Yeast mitochondrial ATPase (to draw up a case study with protein structures already characterised and available in PDB), for which the most negative binding affinities can be found in **Table 4**, along with the corresponding values for the AlphaFold2 structures. LigPlot+ images for the most stable confirmations from the bovine and yeast docked outputs can confirm that enterobactin binds to different residues in the alpha subunit/s of the F1 ATPases as well as neighbouring residues in the beta subunits (**Fig S13-S16**).

**Table 4:** Comparative binding affinity values in kcal/mol for the most stable confirmations for binding with Fe(III)-enterobactin among different organisms.

| ATPase | Most stable confirmation<br>(Fe3+-Ent) | Second most stable confirmation<br>(Fe3+-Ent) |
| --- | --- | --- |
| Human alpha subunit from AlphaFold2 (P25705-ATPA_HUMAN) | -8.6 | -8.4 |
| Bovine whole mitochondrial F1 ATPase (PDB ID 1bmf) | -9.7 | -9.8 |
| Yeast mitochondrial ATPase (PDB ID 2HLD) | -10.2 | -8.9 |

In the case of bovine mitochondrial ATPase, Fe(III)-enterobactin binds to residues of the C (part of the alpha subunit) and D chains (part of beta subunit). In the case of yeast mitochondrial ATPase, the most stable confirmation of Fe(III)-enterobactin binds to the L chain (part of the alpha subunit), and P and M chains (part of the beta subunit), whereas the second most stable confirmation binds to T (part of alpha subunit) and X (part of beta subunit) chains. Docking with RCSB PDB structures yielded more negative values of binding affinity. This extra stability can be due to the role of water molecules, protons (acidic environment) or the surrounding non-alpha subunit residues of the F1 ATPase. For each of the following dockings, interactive protein-ligand diagrams have been obtained in LigPlot+ and PyMol.

### Bovine whole mitochondrial F1 ATPase docking with the most stable conformation of Fe(III)- enterobactin

The interacting residues are in Yellow and the Fe(III)-enterobactin complex is in green (**Fig S13A**). Hydrogen bonding is observed between Fe(III)-enterobactin and Tyr381 and Arg408 residues of the D chain (**Fig S14A**).

### Bovine whole mitochondrial F1 ATPase docking with the second most stable conformation of Fe(III)- enterobactin

As visible in the corresponding LigPlot+ ligand protein interaction diagram (**Fig S14B**), the Fe(III)-enterobactin ligand is in blue in the docking snapshot, to the right is Gln 405 of the chain. To the top left is Tyr381 and to the bottom left is Arg408 of the D chain, all in yellow (**Fig S13B**).

### Yeast mitochondrial ATPase docking with the most stable conformation of Fe(III)-enterobactin

Fe(III)-enterobactin is in blue and the interacting residues are in yellow (**Fig S15A**). The LigPlot+ diagram (**Fig S16A**) shows hydrogen bonding of Fe(III)-enterobactin with Gln332, Asp335 and Arg293 of the L chain, Leu3 and Asn265 of the P chain and Asp319 of the M chain.

### Yeast mitochondrial ATPase docking with Fe(III)-enterobactin of the second most stable conformation

A magnesium atom (green) is present in the vicinity of the ligand-protein interaction site in the docking snapshot in **Fig S15B** (not visible in the 2D protein ligand interaction plot). Fe(III)-enterobactin is in green, and the interacting residues are in yellow. The LigPlot+ diagram (**Fig S16B**) shows the hydrogen bonding of Fe(III)-enterobactin with Ser143 of the T chain and Asp196, Arg192 and Glu193 of the X chain.

### Proposed model of mitochondrial Fe(III) uptake

From the most possible conformations available in **Table 2** for Approaches 2a and 3, it can be suggested that the Fe(II)-enterobactin-alpha complex could be nearly as stable as the complex formed by Fe(III). Although mitochondrial iron uptake and transport has been mostly studied in the context of Fe(II) uptake, it is possible that enterobactin facilitates uptake of Fe(III) in the mitochondria. Once in the mitochondrial matrix, a ferric reductase can reduce the Fe(III) to Fe(II), thus leading to Fe(III) reduction without the requirement of ATP. A Fe(III)-enterobactin complex, can enter the mitochondrial matrix either through the membrane or the stalk of the mitochondrial ATPase.

The import of enterobactin into the mitochondria doesn’t require catalysis of the F0-F1 complex nor an interaction with other subunits, according to Qi and Han (6). It is possible that the Fe(III)-enterobactin complex enters the mitochondrial matrix through either of those two routes and interacts with the alpha subunit without requiring ATP hydrolysis. That should explain why the import of enterobactin does not require catalysis as well as interaction of the other subunits involved. However, as we shall see later, enterobactin does indeed bind to some beta-subunit residues in the ATPase in the case of bovine and yeast ATPases, but that does not refute the case of ATPase catalytic activity being unnecessary for iron uptake. Coming to the model of transport, considering that ATP hydrolysis would not be needed for enterobactin, and hence iron import, the Fe(III)-enterobactin complex must bind to the alpha subunit once it enters the mitochondria. The (Fe(III)-enterobactin)-(alpha) complex is stabilised by a binding affinity of -8.6 and -8.4 kcal/mol as per Approach 4. Once inside the matrix, one of the three events shown in **Fig 1** could occur,

1. Fe(III) is reduced to Fe(II) while still bound to the enterobactin-alpha complex
2. Fe(III)-enterobactin is dissociated from the protein and subsequently the Fe(III) is reduced to Fe(II)
3. Fe(III) is dissociated from the enterobactin-alpha complex and is reduced in the matrix. This last option highly unlikely as it would mean that there is free Fe(III) in the mitochondrial matrix.

**Figure 1:**
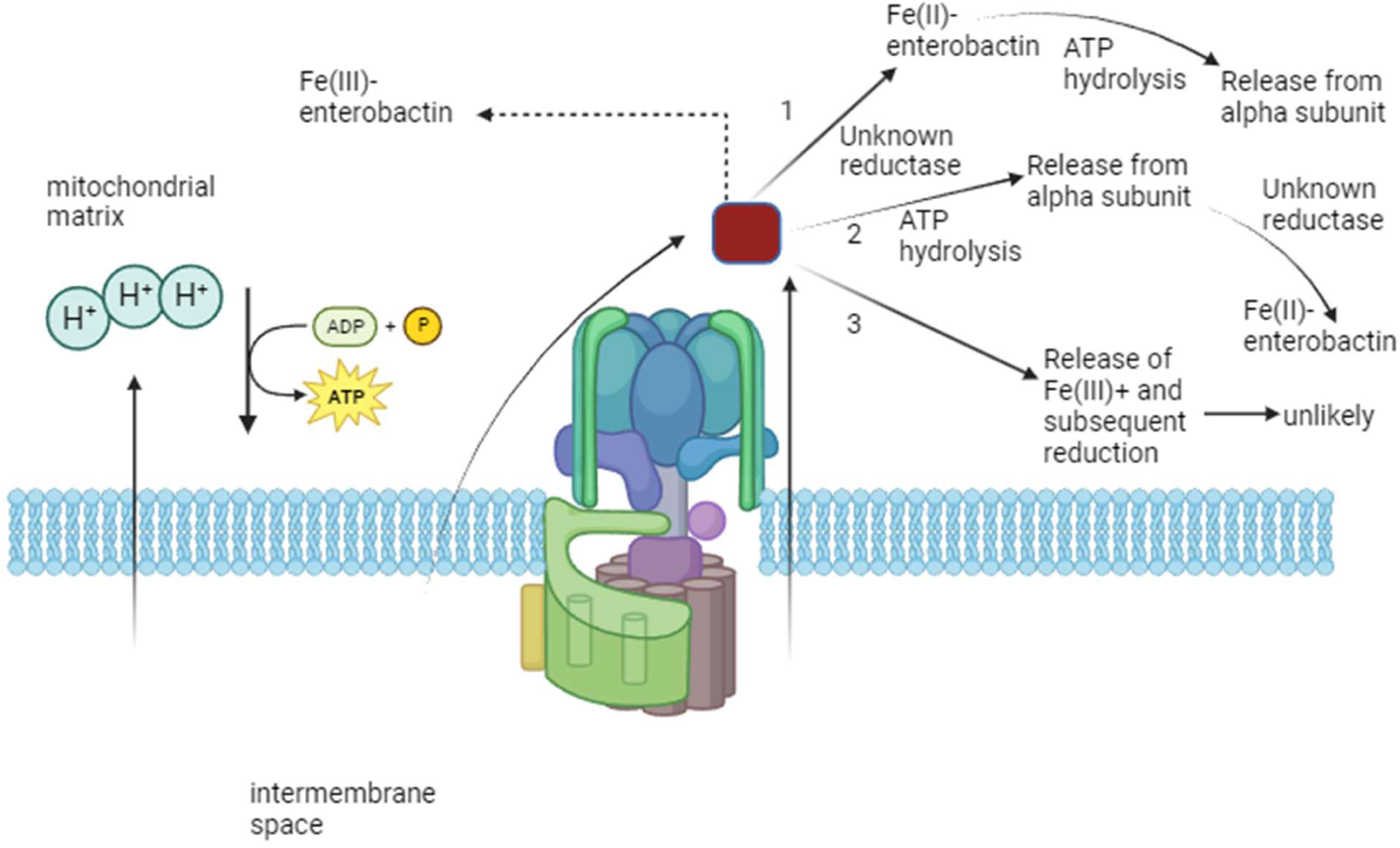
Model for the enterobactin-mediated iron import and reduction in the mitochondria, according to this work. Pathways 1, 2 and 3 are discussed further in the text and differ in the timing of the Fe(III) to Fe(II) reduction and possible ATP hydrolysis requirements. Illustration made in Biorender.

If only the first possibility is considered, the binding affinities in approaches 2a and 2b, i.e., of Fe(II)- enterobactin complexes bound to the alpha subunit, both are more negative than -8.6 and -8.4 (-9.4 and - 8.7 kcal/mol). This implies that the formation of this complex is thermodynamically feasible and would not require any external stabilisation. The formation of Fe(II)-enterobactin-alpha would move the reaction towards the right, which would probably make the reductase reduce more of Fe(III)-enterobactin-alpha to Fe(II)-enterobactin-alpha. This facilitates thermodynamically, if not mechanically, the import of more Fe(III)-enterobactin into the mitochondria. No ATP hydrolysis would be required until this point. To dissociate the Fe(II)-enterobactin-alpha complex into Fe(II)-enterobactin, ATP can be required (for external input of around 9 kcal/mol). One ATP molecule could release around 7-8 kcal/mol. Therefore, it is quite possible that ATP hydrolysis is required for breaking the more stable Fe(II)-enterobactin-alpha complex, once the Fe(III) reduction is completed, leading to Fe(II)-enterobactin.

The binding of enterobactin to the alpha subunit produces a very stable complex and this stability is more than the Fe-enterobactin complex *in silico* (**Table 3**). It can be said that the first pathway of Fe(III) import into the mitochondria is indeed feasible because the second step requires ATP to break the enterobactin-Fe from the alpha subunit. ATP hydrolysis-driven dissociation of the enterobactin-alpha complex might be required for facilitating iron transport. The energetics in this case can offer some support to a possible mechanism of ferric reduction inside the mitochondria, in addition to the already established mechanisms of iron transport. The gradient of binding affinities, from a very unstable Fe-enterobactin complex to a highly stable triple complex of iron, enterobactin and the alpha subunit and eventually dissociation of iron to a slightly less stable complex explains why ATP synthase is not required for importing enterobactin into mitochondria, whereas it would be required for the dissociation of iron inside the mitochondrial matrix.

### Possible candidates for a reductase enzyme

The possibilities for Fe(III) reduction in the mitochondrial matrix required subsequent to the Fe(III) import via enterobactin suggested by this work are depicted in **Fig 2**. The function of frataxin (FXN ) is not clear, but it is involved in assembly of iron-sulphur clusters and has been proposed to act as either an iron chaperone or storage protein (12). It can capture large amounts of Fe(III) and store it in ferric oxide species, which are less reactive than Fe(III) itself. It appears to be able to supply other enzymes with Fe(II). It is possible that FXN can possess the reductase activity required to reduce Fe(III) inside the mitochondria. The ferric chelate reductase FRO3 appears to be present in the mitochondria of plants. Humans appear to have FRO homologues; however, they are presumed to be endosomal/cytosolic.

**Figure 2:**
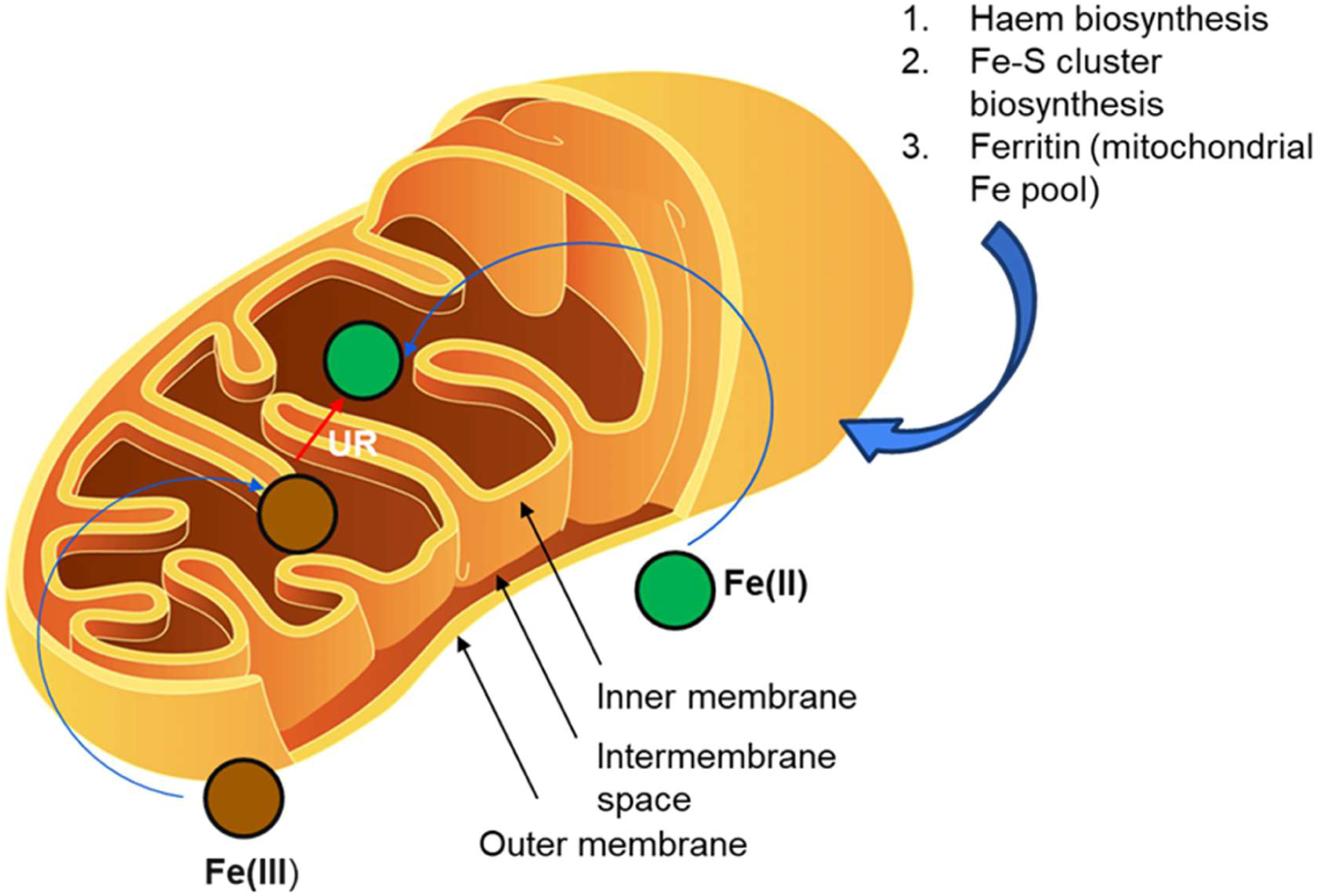
Possible pathways that can reduce Fe(III) to Fe(II) and the fate of Fe(II) in the mitochondria. It is possible that any of the enzymes involved in the three pathways (haem biosynthesis, iron-sulphur cluster synthesis or mitochondrial ferritin can additionally possess reductase activity to fulfil the above model for Fe(III) reduction inside the mitochondria. Illustration made in MS Powerpoint. UR = unknown reductase. The mitochondria drawing is taken from a free online sketch by an unknown author licensed under CC BY-SA-NC.

When iron is absorbed by duodenal enterocytes, cytochrome b reductase 1 (CYBRD1), an iron-regulated ferric reductase, catalyses the reduction of ferric to ferrous ions. Since complex III of the mitochondria in humans contains the CYB gene for cytochrome b (13), it is possible that this enzyme may serve as the required Fe(III) reductase. Indeed, continuous iron reduction in the matrix was proposed in 2000 by Lange et al, who discovered a mitochondrial ferredoxin family protein inducing Fe-S cluster formation in yeast (14). Other matrix ferredoxins are known, such as the mammalian ferredoxins FDX1 and FDX2, which donate electrons to the Fe-S cluster assembly proteins (15). We speculate that FDX1 and FDX2 could be partially at least fulfil the role of electron donors to reduce Fe(III).

All dockings were performed using AutoDock Vina. Required structures were accessed from Pubchem (Fe(II), Fe(III) and Enterobactin structures) as SDF files, AlphaFold2 (Human mitochondrial ATPase alpha subunit) as PDB file and RCSB PDB (Bovine mitochondrial and yeast mitochondrial F1 ATPases) as PDB files. Open Babel GUI (16) was used to convert all file types to AutoDock structure files (PDBQT files). In AutoDock Vina, water molecules associated with proteins were removed and polar hydrogens were added during pre-processing. Missing atoms were checked and repaird and Kollman charges were added to the protein. The appropriate structures were set as ligand and receptor as required in each of the dockings. In all the cases, the larger structure has been set as the receptor and the smaller structure/complex as the ligand. Both ligand and receptor PDBQT files were saved and loaded for docking. The grid coordinates for the receptor were set such that the entire surface of the receptor can be screened for docking sites. The output files were generated and visualised using PyMol. LigPlot+ v.2.2 was licensed by EMBL-EBI for a free academic version and was used to generate ligand-receptor 2D interaction diagrams.

---

**Figure S1:**
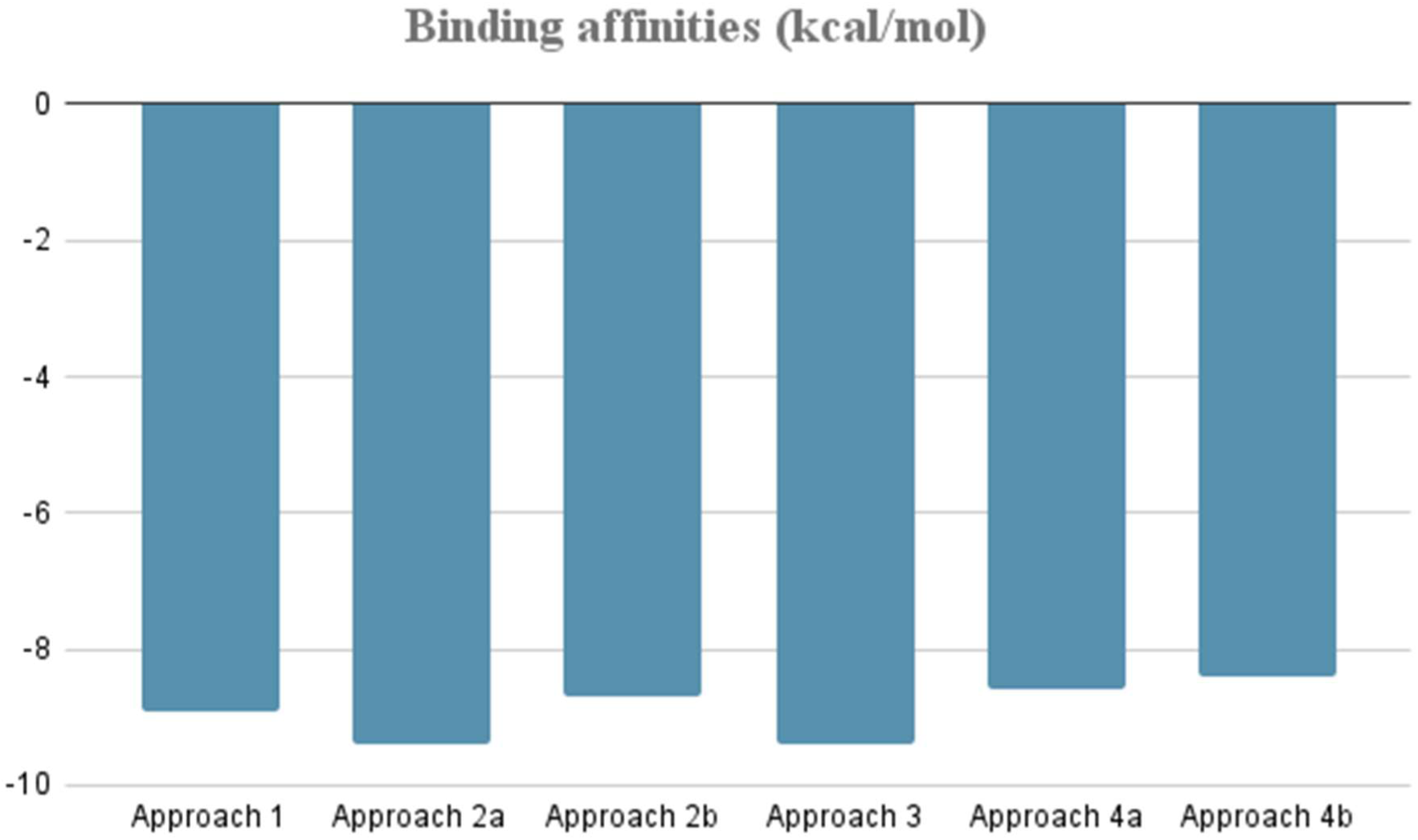
The most negative values of binding affinities obtained from the docking approaches outlined in Table 2.

**Figure S2:**
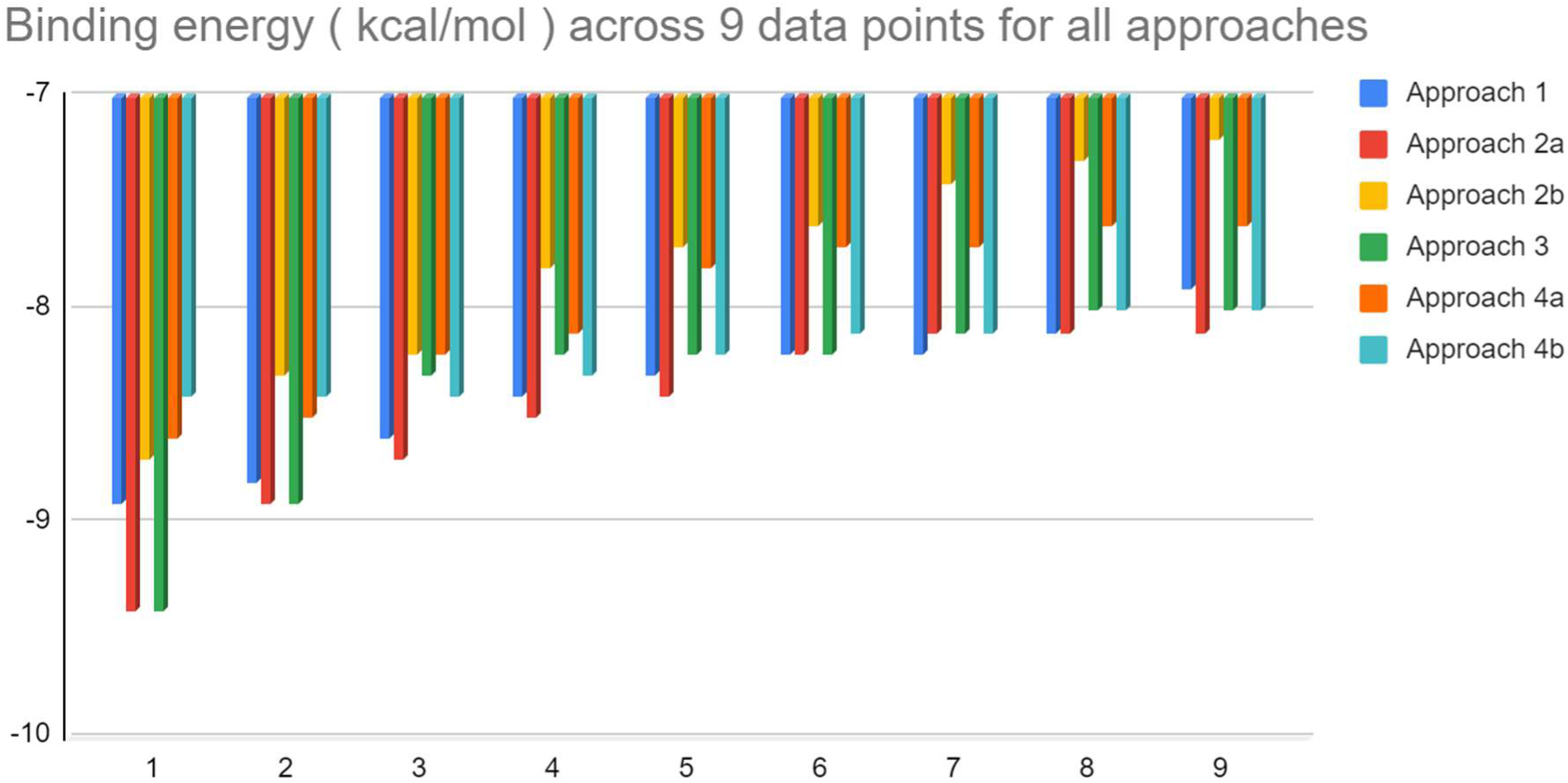
All the values of binding affinities obtained from molecular docking (9 conformations for each approach outlined in Table 1). All the values are tabulated in Table 2. The most negative values for the most stable conformation i.e. the set of points denoted by 1, are of Approach 2a and 3.

**Figure S3:**
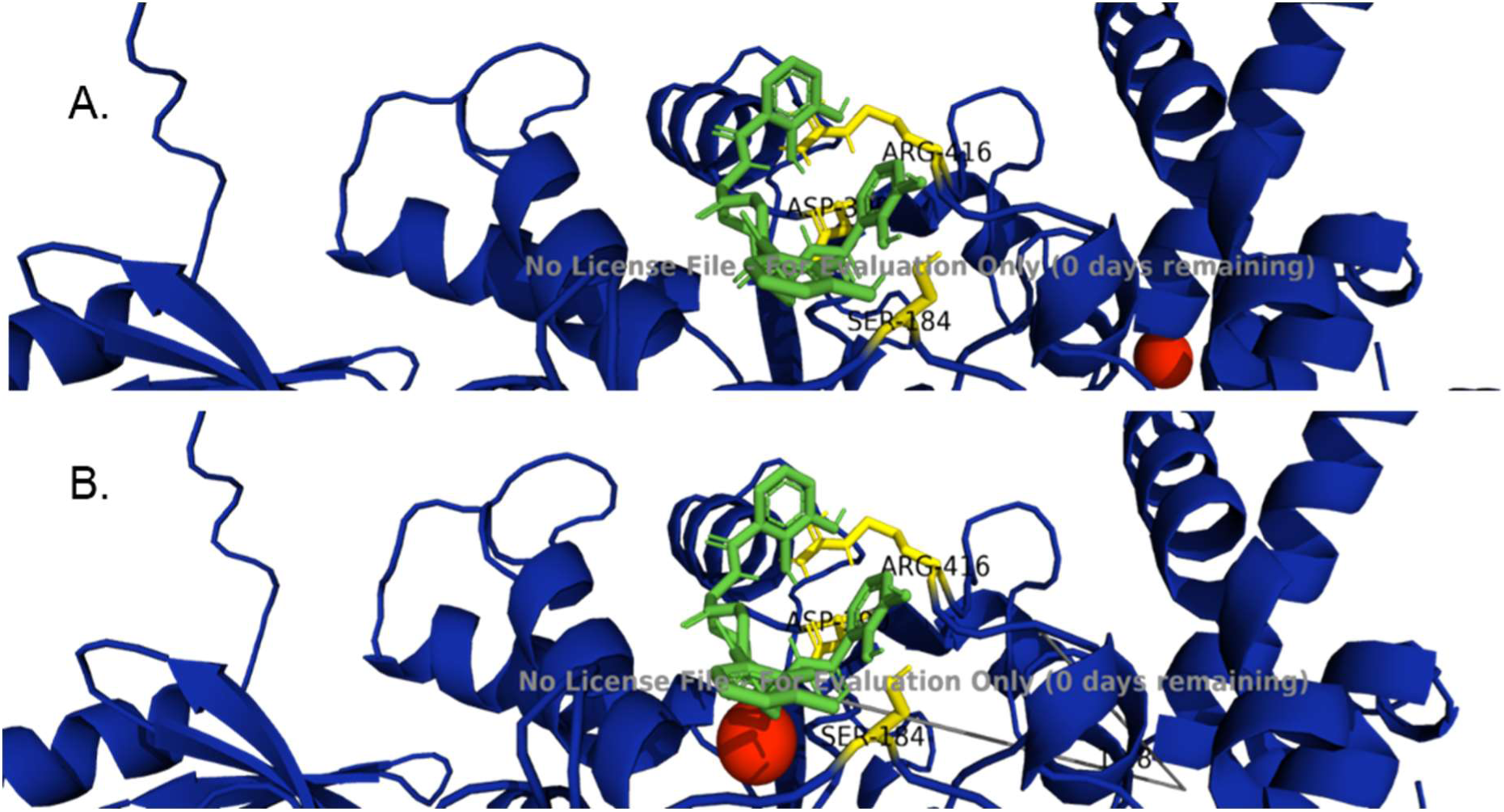
Approach 1 showing the most (A) and second most (B) stable conformations obtained. The red sphere is the Fe(II) ion. The enterobactin is shown in green. The interacting residues are shown in yellow (Arg 416, Ser 184 and Asp 390). Detailed molecular interaction is present in Figure S7.

**Figure S4:**
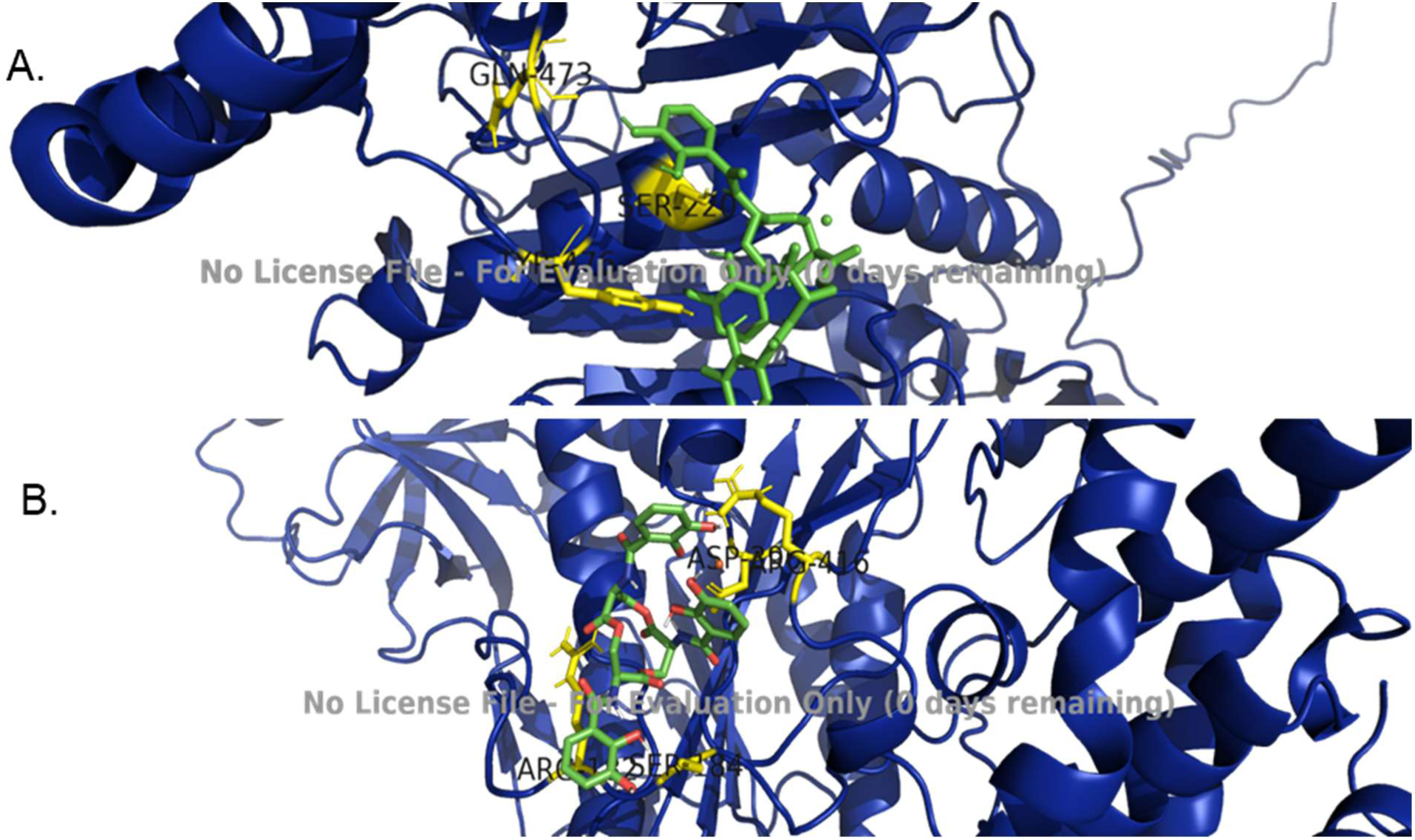
(A) Approach 2a showing the most stable conformation obtained after the second docking. Enterobactin is shown in green and the small green sphere along the enterobactin ring is the Fe(II) ion. The interacting residues of the alpha subunit are shown in yellow. (B) The most stable conformation obtained after the second docking for Approach 2b. The interacting residues are shown in yellow (For 2a, enterobactin is seen to interact with the Ser220, Gln473 and Tyr476 residues of the chain A of the alpha subunit of the mitochondrial ATPase, while for 2b the residues are Arg182, Ser184, Asp390 and Arg416). Detailed molecular interactions are presented in Figures S8 and S9.

**Figure S5:**
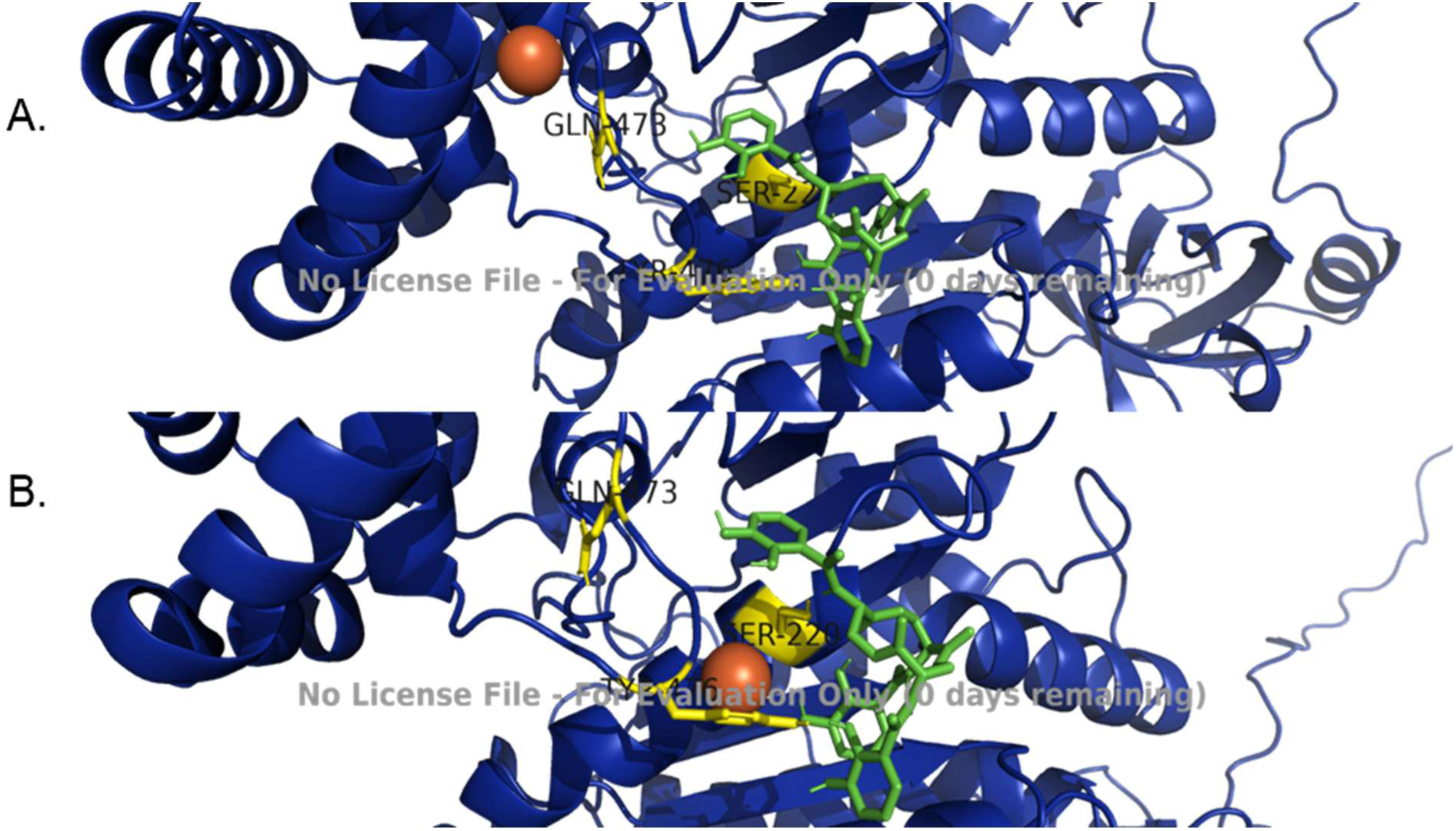
Approach 3 showing the most stable conformation in (A). In this approach however, the sixth most stable conformation (B) yields the closest spatial arrangement of the Fe(III) ion with the enterobactin and the binding residues of the alpha subunit. The red sphere is the Fe(III) ion. The enterobactin is shown in green. The interacting residues are shown in yellow. Enterobactin interacts with Gln473, Ser220 and Tyr476 residues of chain A of the alpha subunit of mitochondrial ATPase by hydrogen bonds). Detailed molecular interactions are presented in Figure S10. Note: In this approach, however, the 6th most stable conformation (B) yields the closest spatial arrangement of the Fe(III) ion with the enterobactin and the binding residues of the alpha subunit. Panel A shows the lowest energy in docking, but in terms of proximity to the residues that are shown to be involved in interaction using enterobactin as ligand, the lower picture B (which is the 6th conformation) shows the ferric ion nearest to the enterobactin(green)-alpha subunit (yellow) residue complex. This implies that in sillico, without considering the proton rich environment of the mitochondria and other associated environmental factors, the energetically most stable confirmation is not necessarily the one where all the three components [Fe(III), alpha subunit and Enterobactin] are in the closest proximity. This might mean that additional environmental factors can potentially stabilise the less stable structure such that it becomes more or less stable as the most stable confirmation in vivo, or that the most stable confirmation itself involves some mediating molecule that makes it possible for Enterobactin to functionally interact with the Fe(III) else the iron atom being far from enterobactin does not seem functionally feasible in vivo.

**Figure S6:**
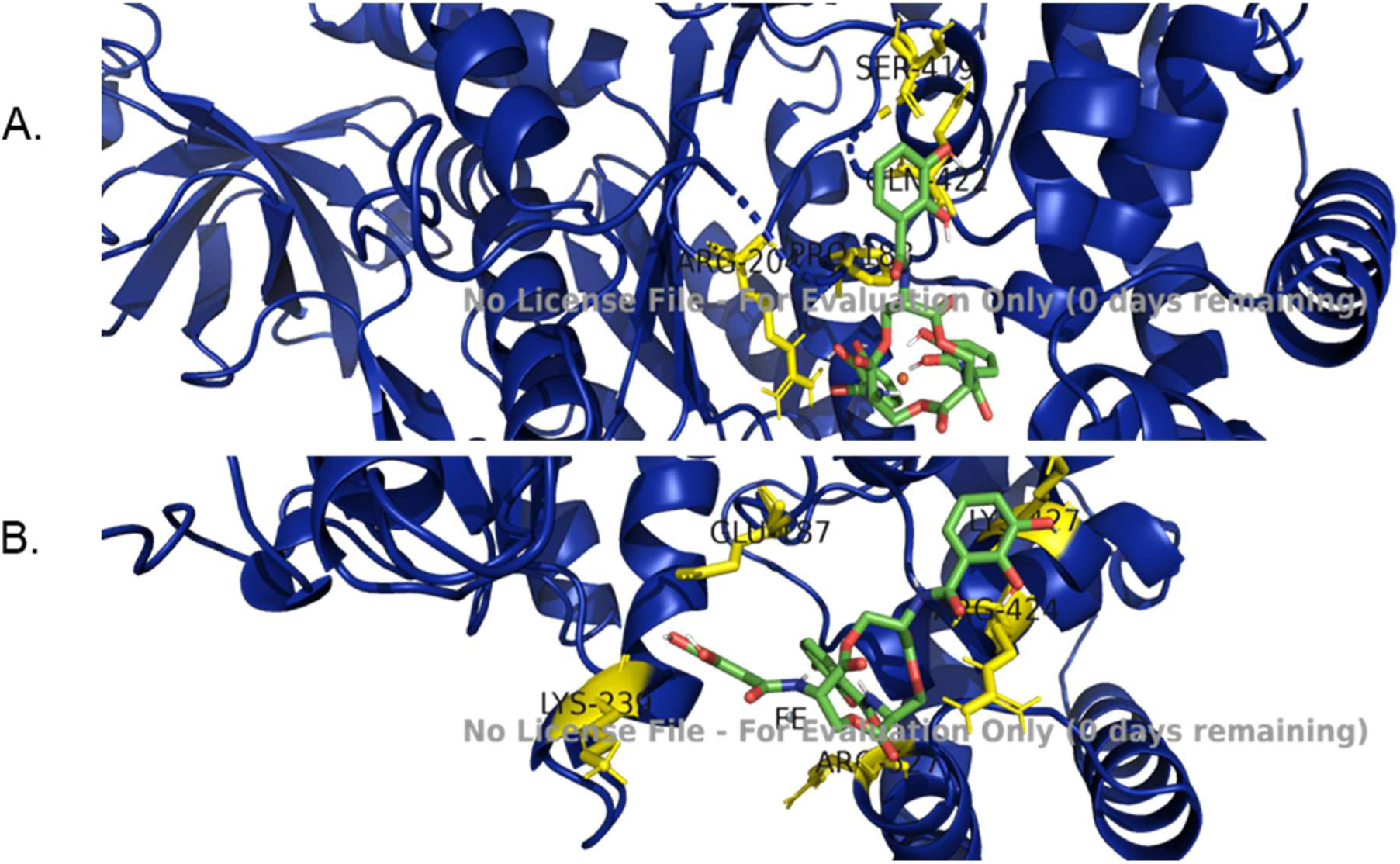
(A)The most stable conformation obtained after the second docking for Approach 4a. (B) Approach 4b showing the most stable conformation obtained after the second docking. Fe(III) is labelled as FE in (B). The interacting residues are shown in yellow and enterobactin is in green. For 4a, enterobactin is seen to interact with the Arg204, Gln422, Ser419, Pro188 residues of the chain A of the alpha subunit of the mitochondrial ATPase by hydrogen bonds, while for 2b the residues are Lys239, Gln187, Arg424, Arg527 and Lys427. Detailed molecular interactions are presented in Figures S11 and S12.

**Figure S7:**
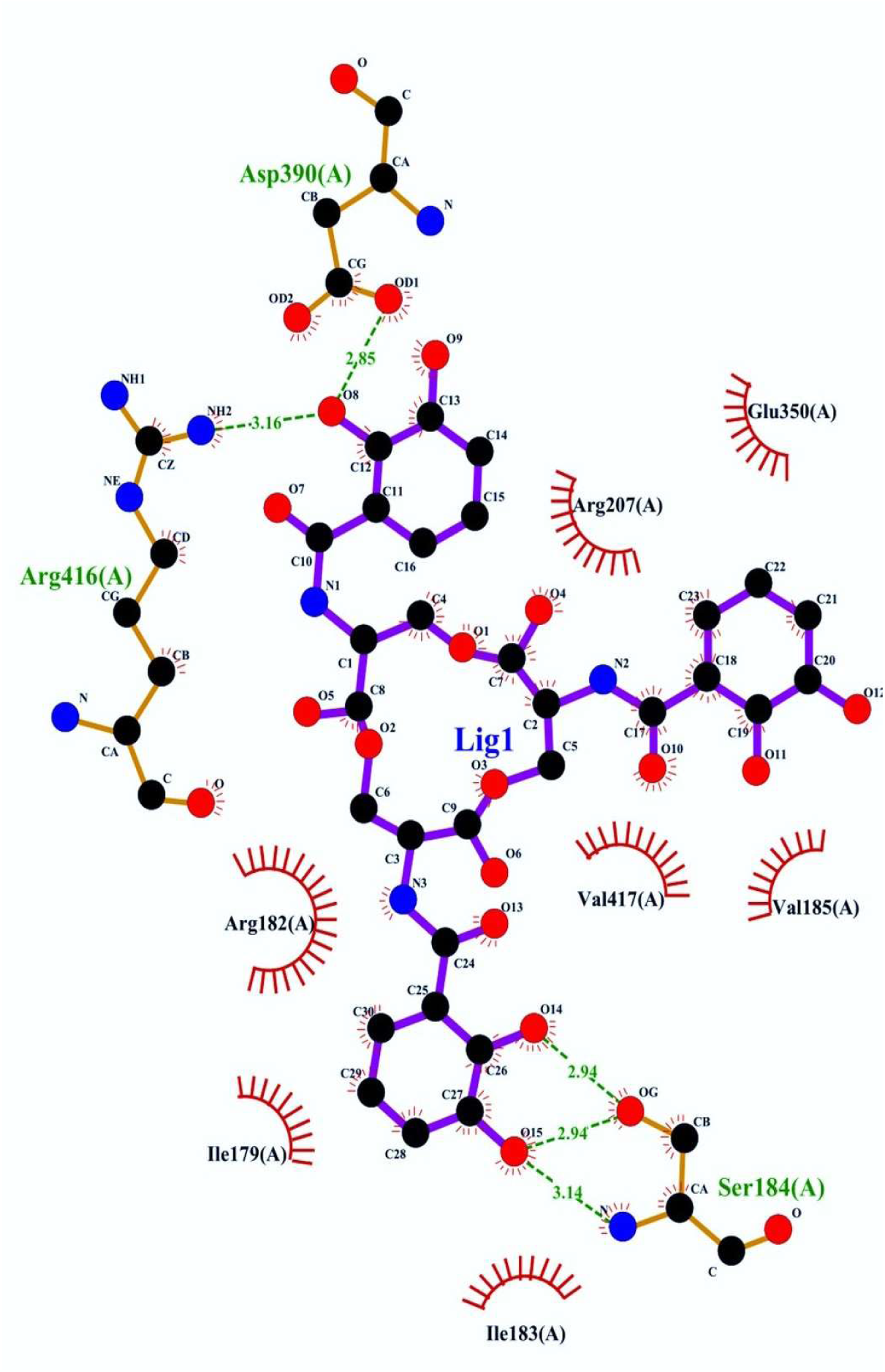
LIGPLOT+ plot of approach 1. Green represents hydrogen bonds, while red arcs show covalent interactions. Solid lines show covalent bonds.

**Figure S8:**
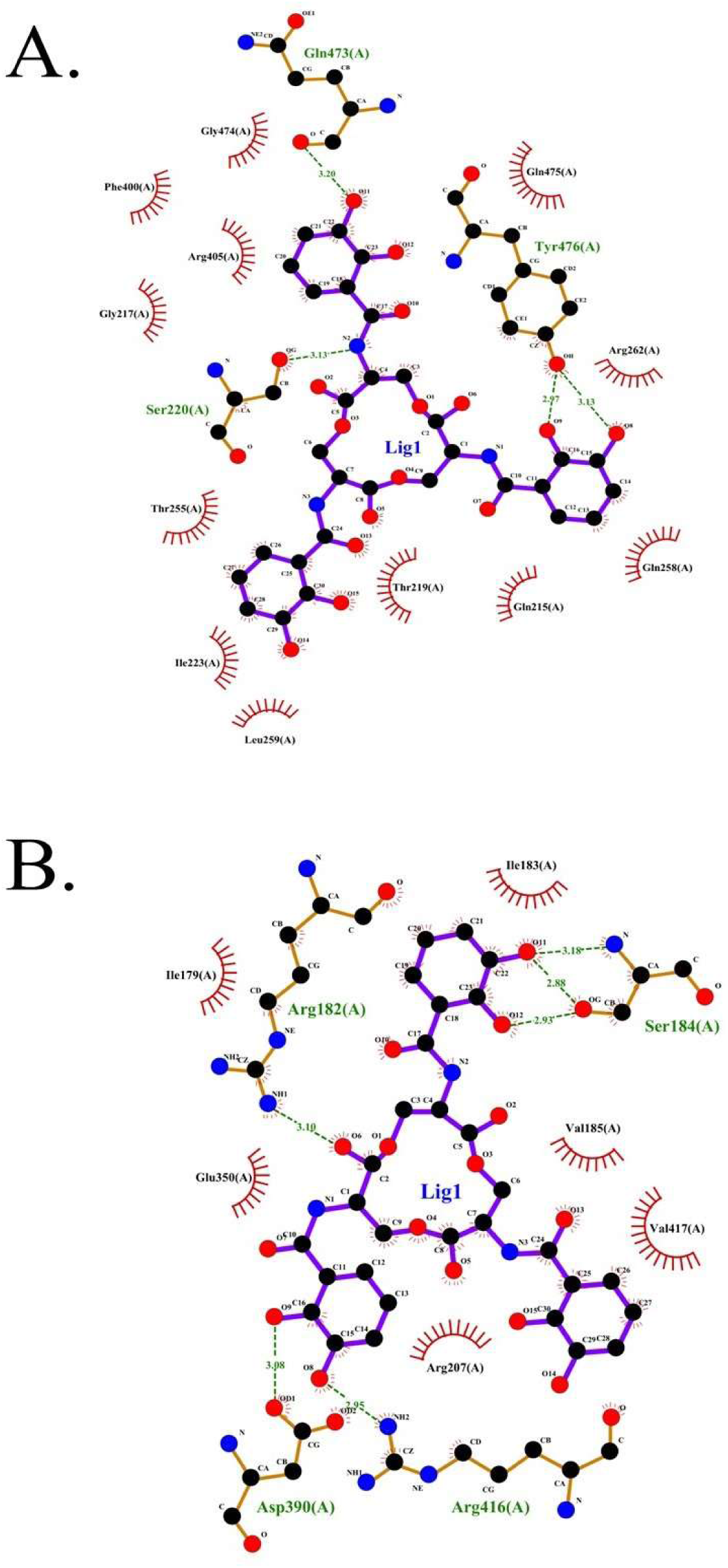
LIGPLOT+ plots of approach 2a (A) and 2b (B). For 2a, enterobactin is seen to interact with the Ser220, Gln473 and Tyr476 residues of the chain A of the alpha subunit of the mitochondrial ATPase by hydrogen bonds, while for 2b the residues are Arg182, Ser184, Asp390 and Arg416, detailing the docking snapshot in Figure S4. Green represents hydrogen bonds, while red arcs show covalent interactions. Solid lines indicate covalent bonds.

**Figure S9:**
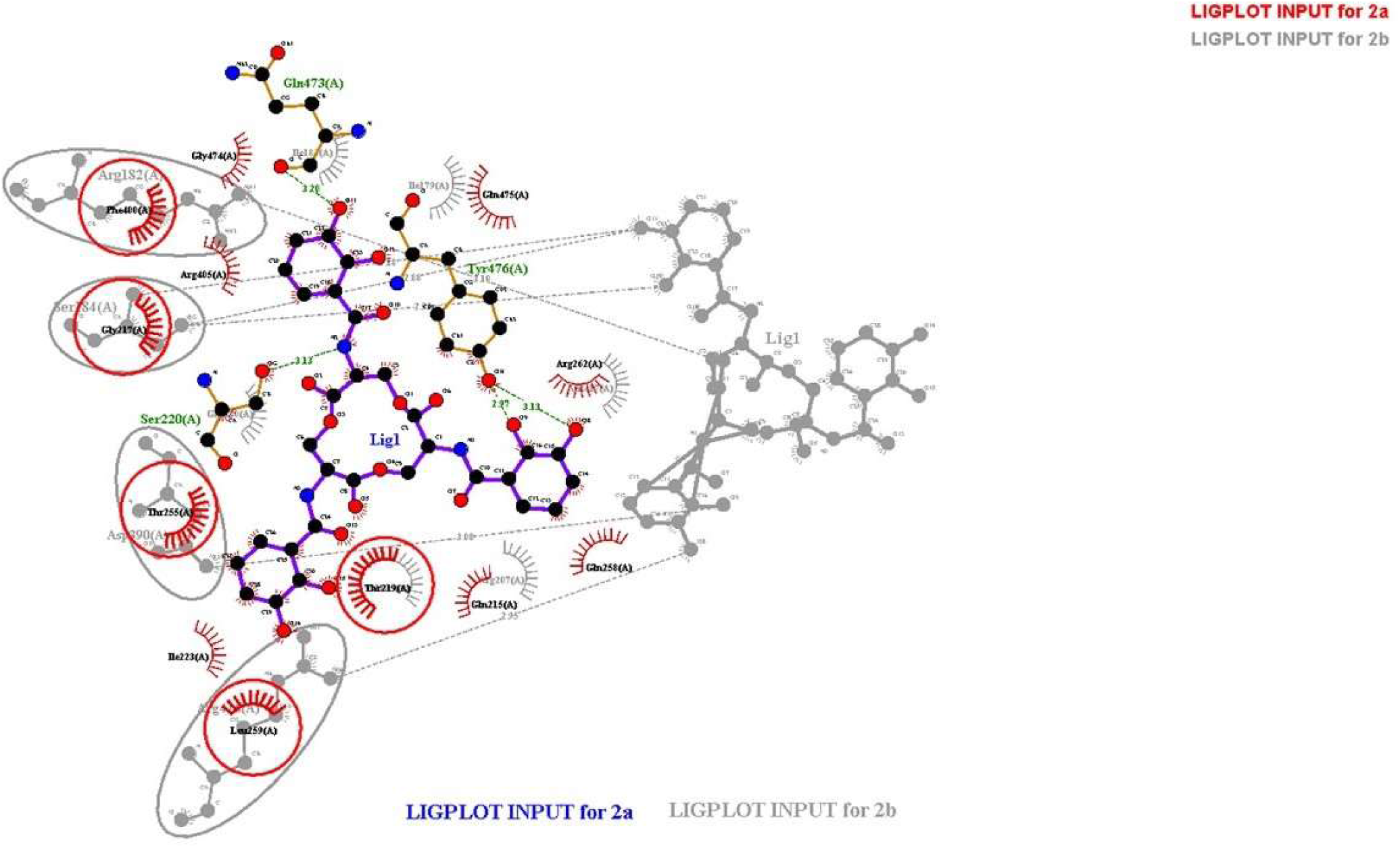
Overlap of LIGPLOT+ plots of Approaches 2a (red) and 2b (grey). The two previous LigPlot+ diagarams overlap to highlight the different amino acid residues involved in ligand-receptor interaction between approaches 2a and 2b.

**Figure S10:**
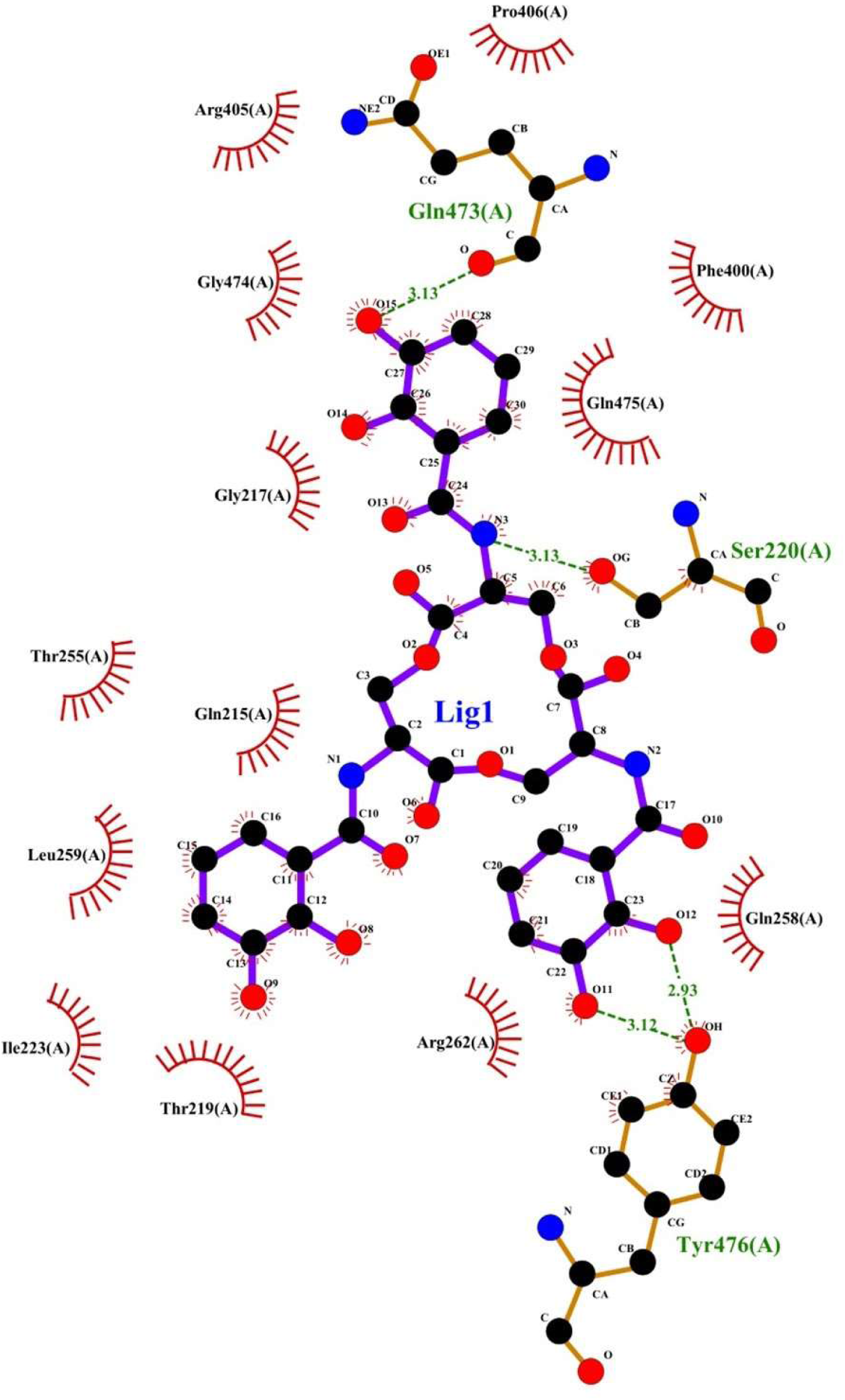
LIGPLOT+ plot of Approach 3. Detailed protein interaction diagram for Approach 3, showing that Enterobactin interacts with Gln473, Ser220 and Tyr476 residues of chain A of the alpha subunit of mitochondrial ATPase by hydrogen bonds. Green represents hydrogen bonds, while red arcs show covalent interactions. Solid lines show covalent bonds.

**Figure S11:**
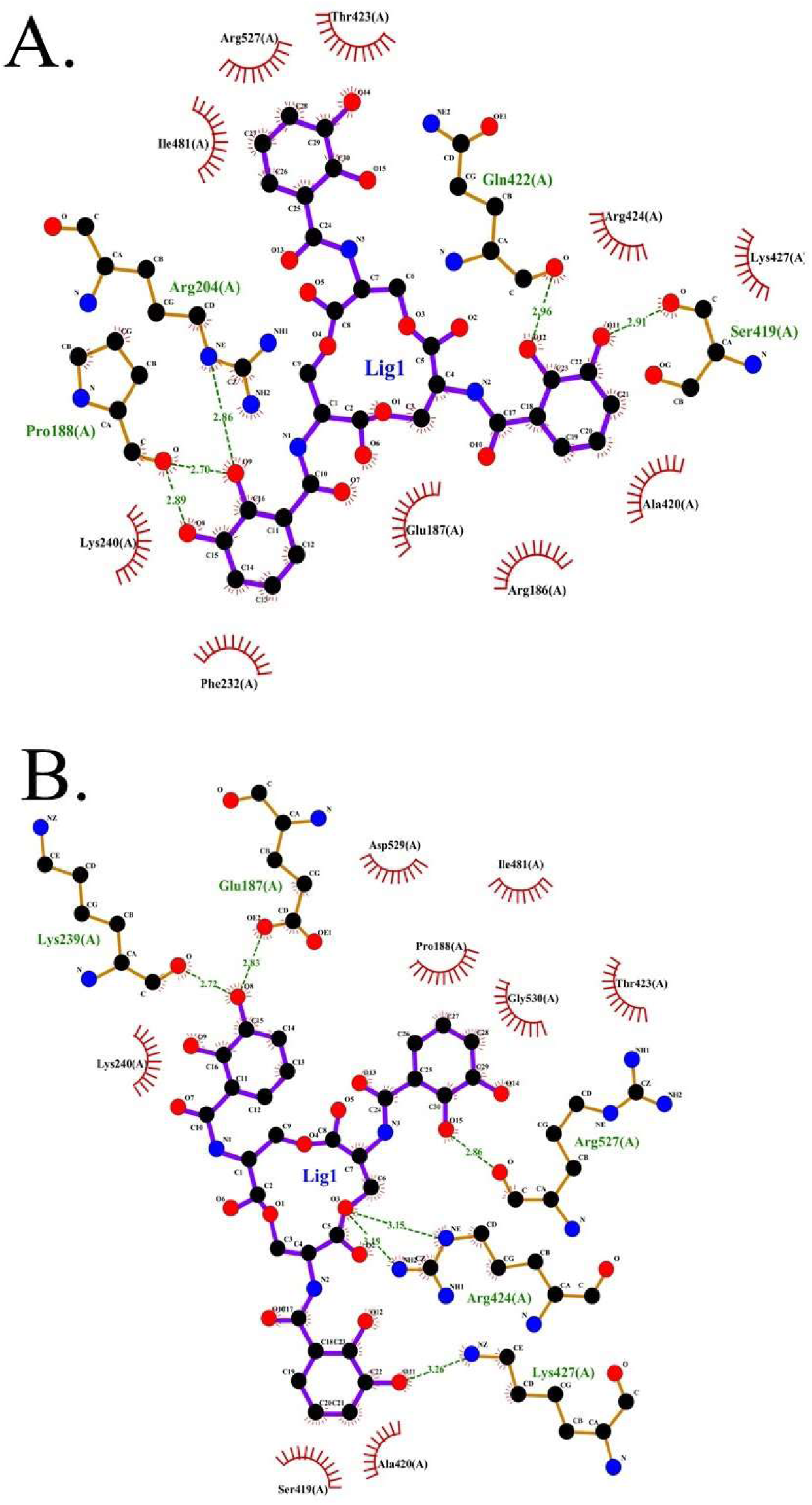
LIGPLOT+ plots of approach 4a (A) and 4b (B). Interactive diagrams for approaches 4a and 4b. For 4a, enterobactin is seen to interact with the Arg204, Gln422, Ser419, Pro188 residues of the chain A of the alpha subunit of the mitochondrial ATPase by hydrogen bonds, while for 2b the residues are Lys239, Gln187, Arg424, Arg527 and Lys427, detailing the docking snapshot in Figure S6. Green represents hydrogen bonds, while red arcs show covalent interactions. Solid lines show covalent bonds.

**Figure S12:**
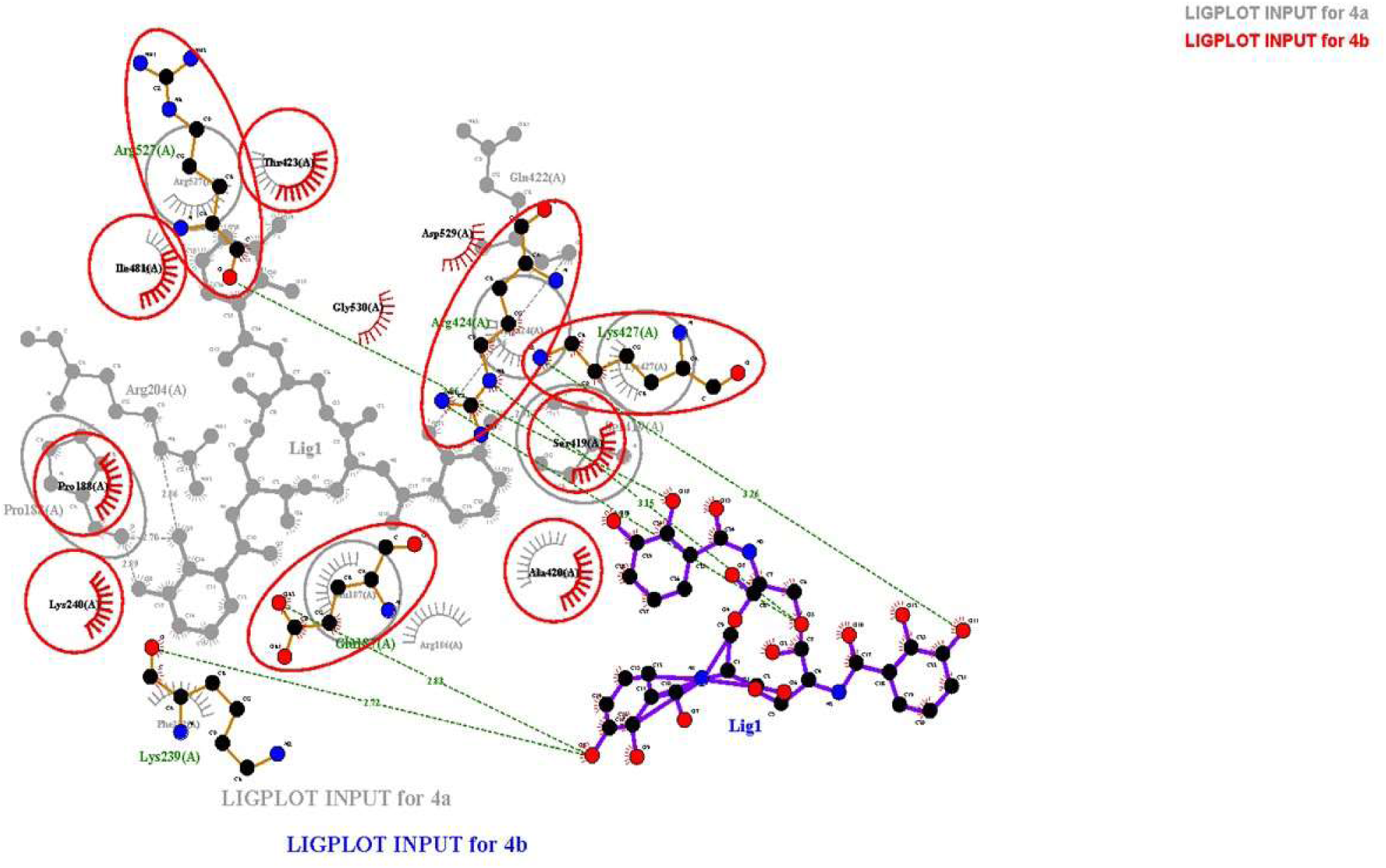
Overlap of LIGPLOT+ plots of approach 4a (grey) and 4b (red). The two previous LigPlot+ diagrams overlap to highlight the different and common amino acid residues involved in ligand-receptor interaction between approaches 4a and 4b.

**Figure S13:**
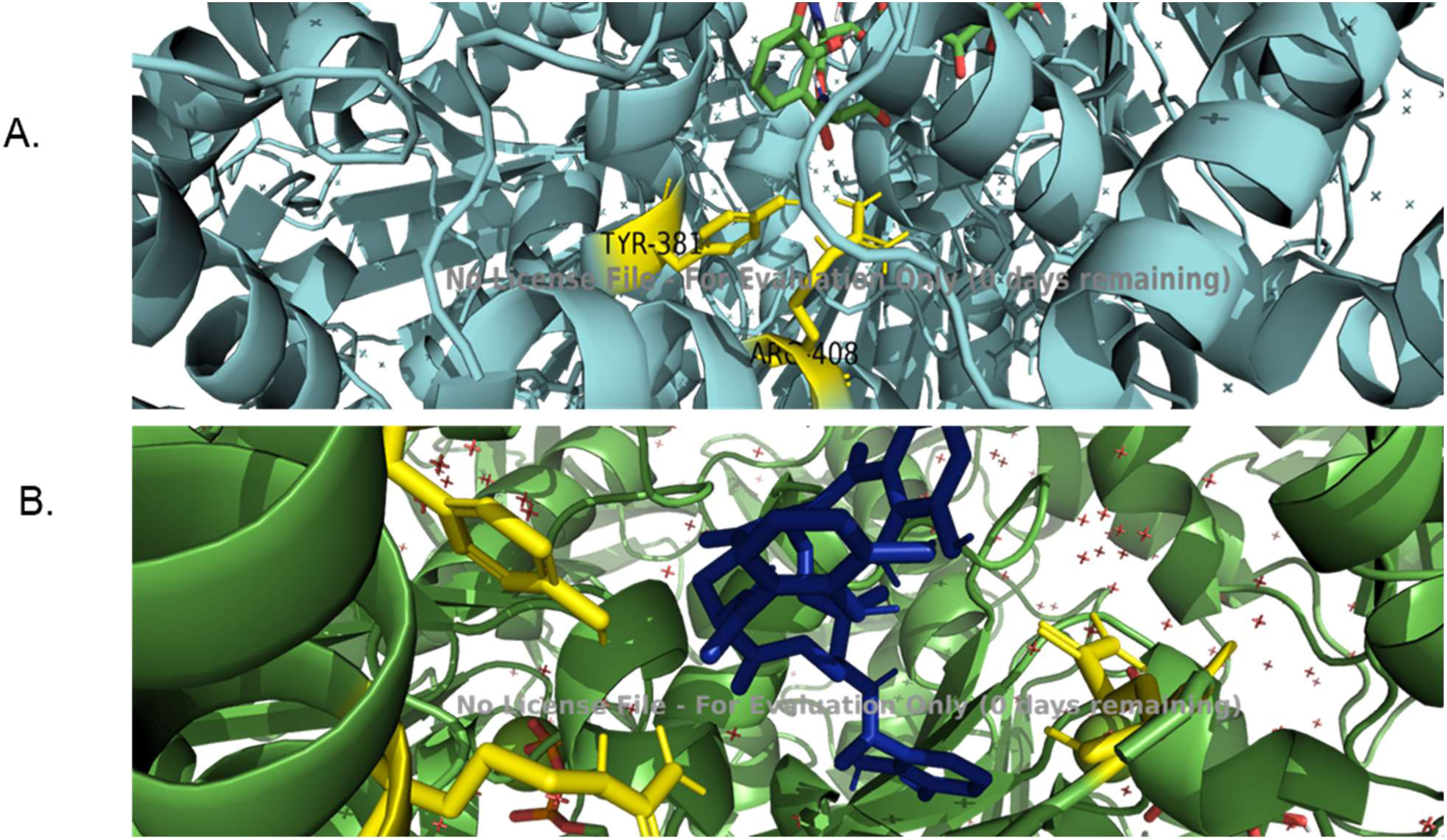
(A) Most stable conformation of bovine whole mitochondrial F1 ATPase docking with Fe(III)-enterobactin. (B) Second-most stable conformation of bovine whole mitochondrial F1 ATPase docking with Fe(III)-enterobactin. The interacting residues are in Yellow (Tyr381 and Arg408 for A and Gln405, Tyr381 and Arg408 for B). The Fe(III)-enterobactin complex is shown in green in panel A and blue in panel B.

**Figure S14:**
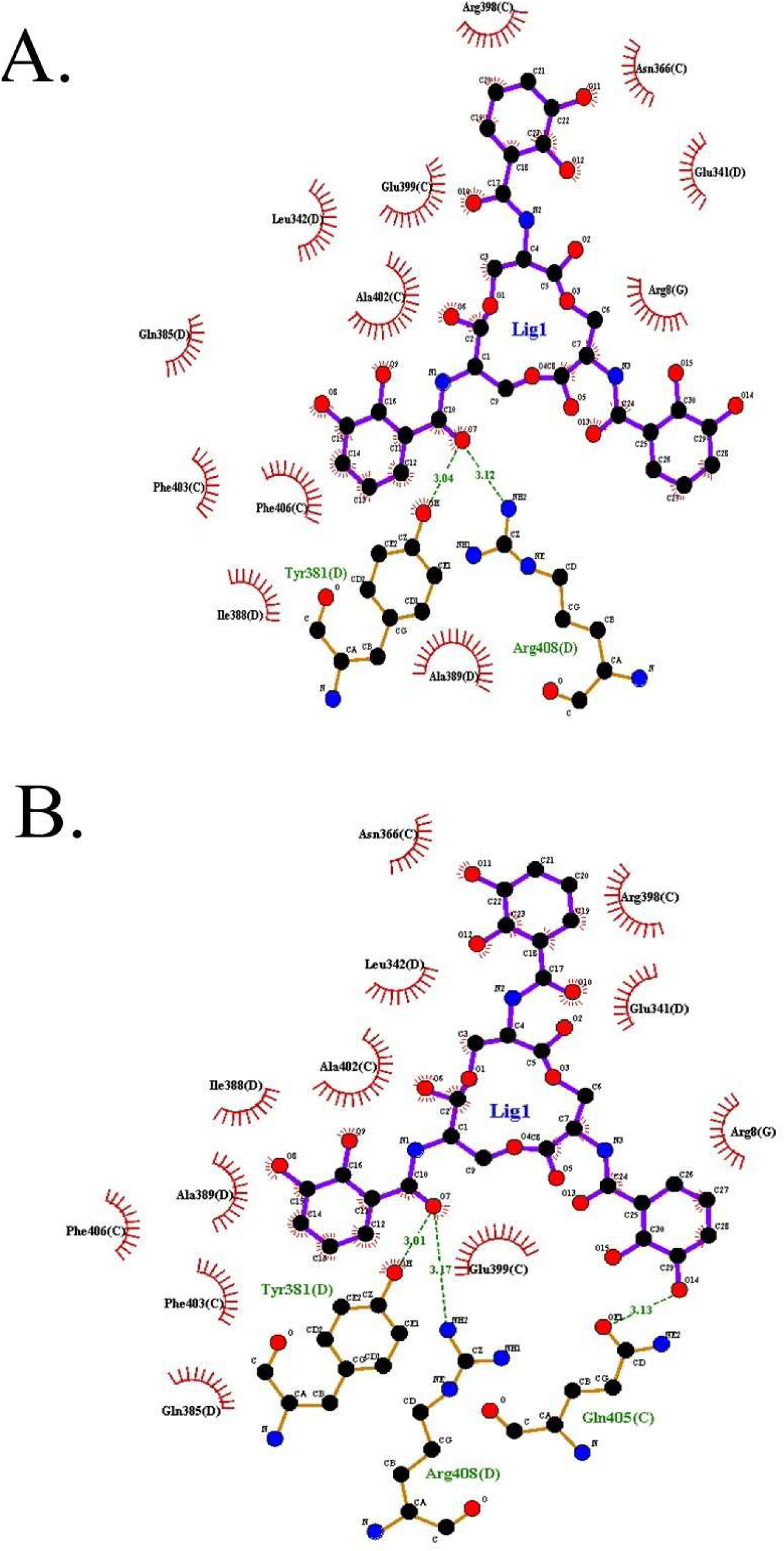
LIGPLOT+ plots for (A) Most stable confirmation of bovine whole mitochondrial F1 ATPase docking with Fe(III)-enterobactin and (B) Second-most stable confirmation of bovine whole mitochondrial F1 ATPase docking with Fe(III)-enterobactin.

**Figure S15:**
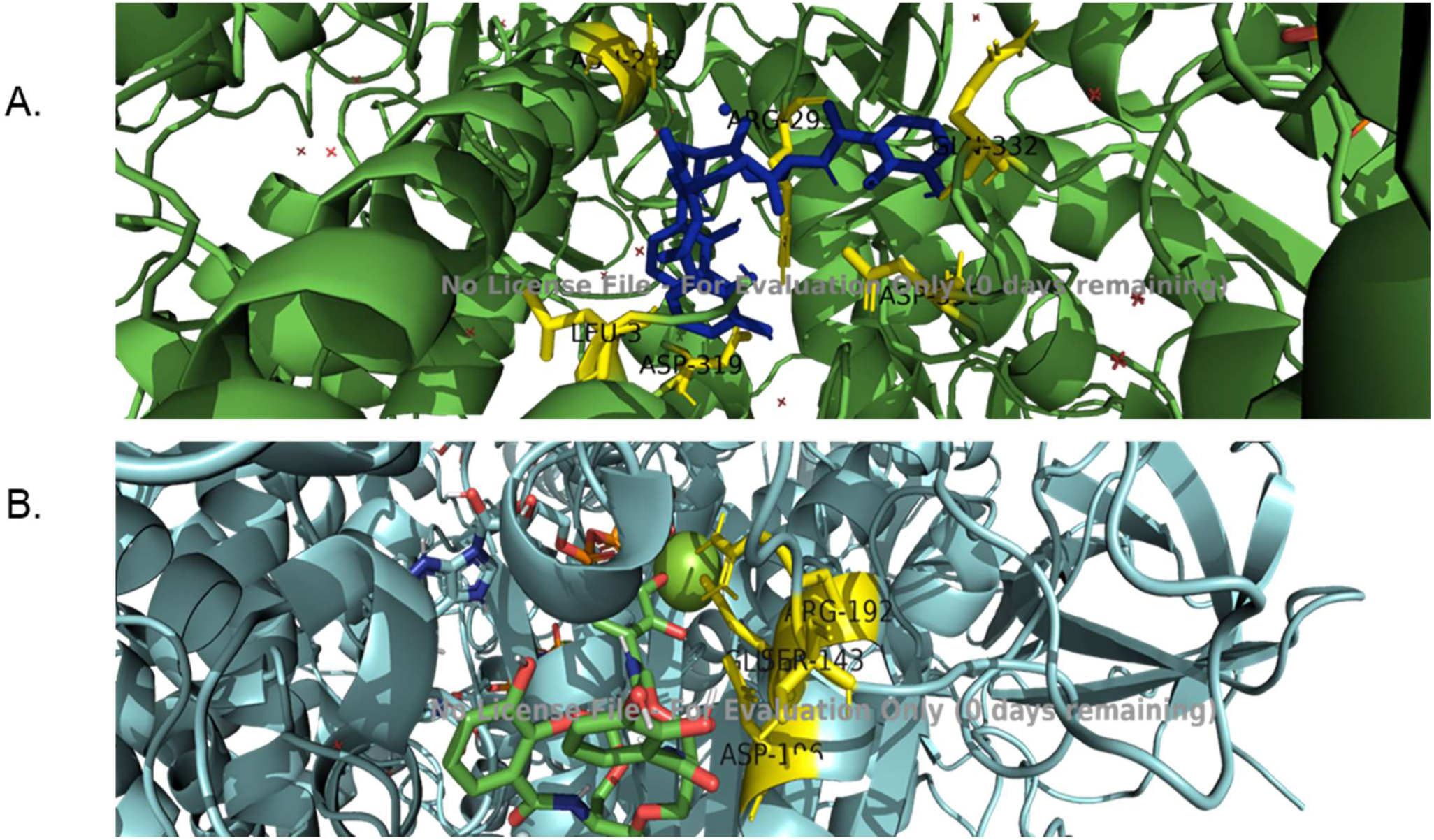
(A) Most stable confirmation of yeast mitochondrial ATPase docking with Fe(III)-enterobactin. (B) Second-most stable confirmation of yeast mitochondrial ATPase docking with Fe(III)-enterobactin. The interacting residues are in Yellow (Gln332, Asp335 and Arg293 of the L chain, Leu3 and Asn265 of the P chain and Asp319 of the M chain for A. For B, the residues are Ser143 of the T chain and Asp196, Arg192 and Glu193 of the X chain. The Fe(III)-enterobactin complex is in blue in A and green in B.

**Figure S16:**
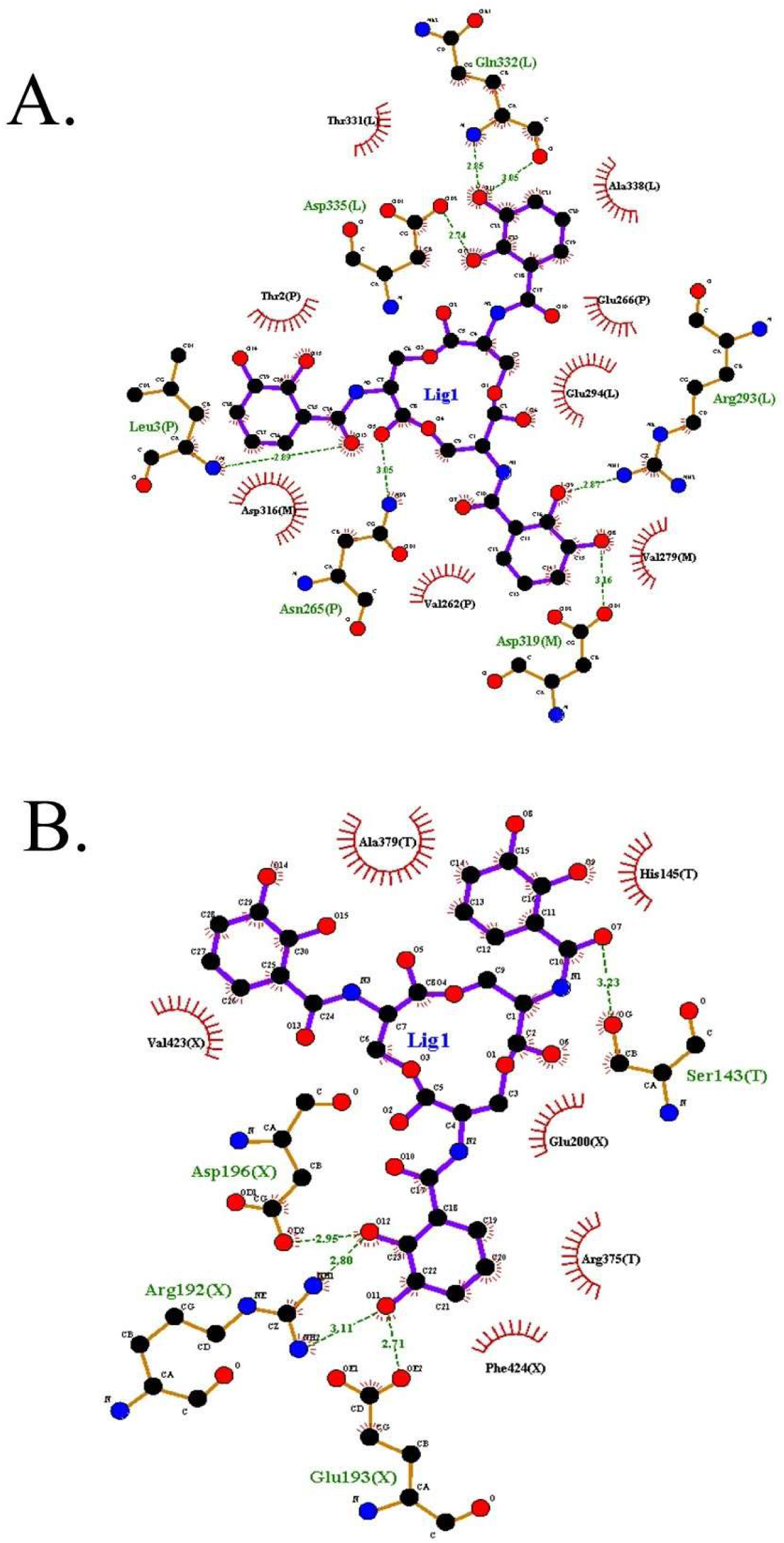
LIGPLOT+ plots for (A)Most stable confirmation of yeast mitochondrial ATPase docking with Fe(III)-enterobactin and (B) Second-most stable confirmation of yeast mitochondrial ATPase docking with Fe(III)-enterobactin.

